# talklr uncovers ligand-receptor mediated intercellular crosstalk

**DOI:** 10.1101/2020.02.01.930602

**Authors:** Yuliang Wang

## Abstract

Single cell RNA-seq measures the transcriptomes of many cell types across diverse conditions. However, an emerging challenge is to uncover how different cell types communicate with each other to maintain tissue homeostasis, and how inter-cellular communications are perturbed in diseases. To address this problem, we developed talklr, an information theory-based approach to uncover potential ligand-receptor interactions involved in tissue homeostasis and diseases. Compared to existing approaches that analyze changes in each gene in each cell type separately, talklr uses a holistic approach to simultaneously consider expression changes in both ligands and receptors across multiple cell types and conditions. talklr outperformed existing approaches in identifying ligand-receptor interactions, including those known to be important for tissue-specific functions and diseases across diverse datasets. talklr can reveal important signaling events in many biological problems in an unbiased way, and will be a valuable tool in single cell RNA-seq analysis. talklr is available as both an interactive website and an R package.

## Introduction

Single cell transcriptomics has become a popular tool to reveal cellular heterogeneity and dynamics in many biological problems^1–3^. Common uses are to uncover novel cell types and subtypes, chart cell state transitions, identify cell type-specific marker genes, and find cell type-specific differentially expressed genes in response to perturbations (e.g., developmental cues, genetic manipulation, diseases, drug treatments)^4^. Increasingly, many studies map single cell transcriptomics data onto curated ligand-receptor interaction networks^5^ to identify potential inter-cellular signaling between cell types in one condition, and how such inter-cellular signaling networks changes across conditions^6–9^.

There are three existing approaches that map single cell transcriptomics data onto ligand-receptor signaling networks. In the first approach, when a ligand is expressed in one cell type and the cognate receptor is expressed in a second cell type, a connection is drawn between this pair of cell types. This is followed by counting how many co-detected ligand-receptor gene pairs exist between each pair of cell types, and the number of cell-cell interactions each ligand-receptor gene pair participates in^6, 10^. In the second approach, when both the ligand and the receptor are specifically enriched in one cell type (not necessarily the same cell type), a connection is drawn between this pair of cell types ^7, 8, 11^. The third approach first quantifies ligand-receptor interactions by multiplying the expression level of the ligand in cell type *i* with the expression of the cognate receptor in cell type *j* to obtain an interaction score, and then prioritizes the ligand-receptor gene pairs with large interaction scores^9, 12^.

Although very valuable at revealing interesting inter-cellular communication *in vivo*, existing approaches face several problems. Firstly, existing approaches do not quantitatively evaluate the distribution of ligand-receptor expression specificity in a continuous way. They either ignore expression specificity (as long as a gene is expressed above a user-defined threshold), or require a hard cell type-specific enrichment cutoff (e.g., fold enrichment >2, false discovery rate<0.05). The latter approach assumes a one-to-one cell-cell interaction for any ligand-receptor pair, which may be too restrictive in many real biological systems. We will show that selecting cell type-specific marker genes is a sufficient but not necessary condition for the ligand-receptor gene pair to exhibit cell-cell interaction specificity – there are many ways for a ligand-receptor interaction to be highly specific without both genes being uniquely expressed in one cell type (e.g., **Figure 1**). Secondly, existing approaches perform univariate analysis for both the genes and cell types, analyzing expression changes for each gene in each cell type independently. However, the biological relevance of a gene’s expression change in one cell type may depend on its baseline expression level in another cell type. For example, a receptor’s statistically significant expression increase from 2 FPKM to 10 FPKM in cell type A may not be very meaningful when its baseline expression level in cell type B is 200 FPKM.

**Figure 1.**
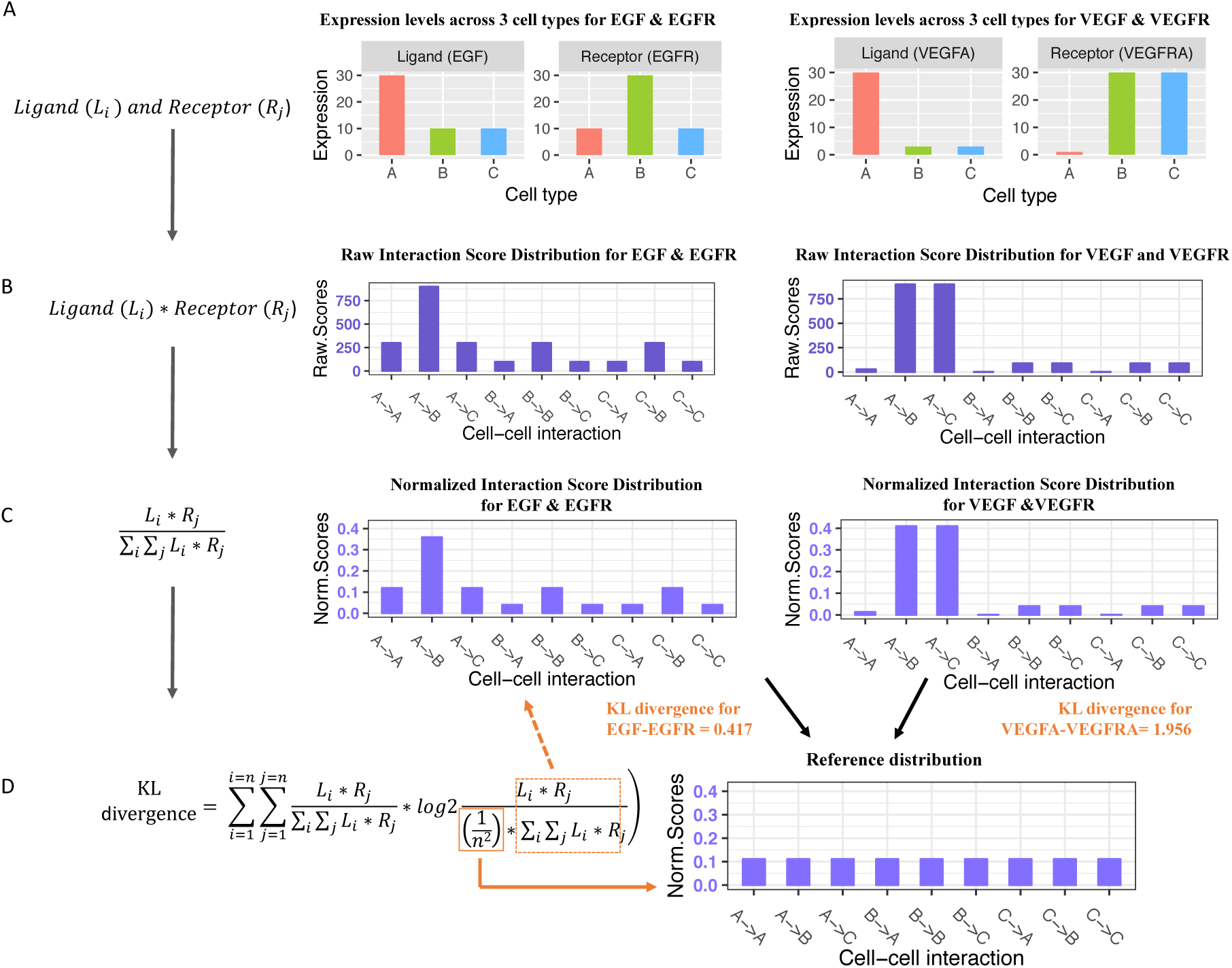
Workflow of talklr in single condition mode using two toy examples. The expression values shown in A are entirely hypothetical; values in B,C,D are derived from them exactly as the talklr method would do. **A.** Expression level of ligands and receptors in three cell types A, B, and C. For EGF-EGFR (left panel) and VEGF-VEGFR (right panel). **B.** Distribution of raw interaction scores for EGF-EGFR and VEGF-VEGFR. Interaction score is the product of ligand expression in cell type i (sender) and receptor expression in cell type j (receiver) **C.** Distribution of normalized interaction scores. Raw interaction scores are normalized by dividing the sum of interaction scores across all pairwise cell-cell interactions. **D.** Kullback-Leibler (KL) divergence quantifies the differences between the observed distribution of normalized interaction scores for a ligand-receptor pair and the reference distribution (1/n^2^ for each of n^2^ cell-cell interactions), which is 0.417 and 1.956 for EGF-EGFR and VEGF-VEGFR respectively. Larger KL divergence in this case means that distribution of interaction scores is concentrated in a few cell-cell interactions, while other cell-cell interactions receive low scores. Note that VEGF-VEGFR achieved higher KL divergence even though the receptor VEGFR is not cell type-specific.

To overcome these problems, we adapted Kullback-Leibler (KL) divergence from information theory to develop the talklr method (intercellular cross****talk**** using ***l*** igand-***r*** eceptor). KL divergence quantifies the difference between one probability distribution and a second, reference probability distribution. The key innovation of talklr that it examines the entire interaction score distribution instead of individual gene expression values. We show that talklr can quantify the expression specificity of ligand-receptor pairs and uncover inter-cellular interactions enriched for known tissue-specific functions across many datasets (e.g., the Tabula Muris Project of 20 mammalian tissues^13^). Furthermore, we demonstrate that talklr can identify potential ligand-receptor signaling rewiring among interacting cell types in disease, development, and aging. talklr outperforms commonly used workflows in both tasks. The advantages of talklr are that 1) it is a quantitative method that avoids arbitrary cutoffs, 2) it considers the entire expression distribution of both ligand and receptor genes across multiple cell types as opposed to univariate analysis of genes and cell types, and 3) when comparing two conditions, it also considers the expression change of one gene in one cell type in the context of the gene’s baseline expression and expression changes in other cell types. The talklr method will be a valuable addition to standard single cell transcriptomics analysis workflows. We developed an R package for talklr so that it can be easily integrated into existing single cell RNA-seq workflow. We also developed an interactive website to enable researchers without programming experiences to perform explorative analysis and visualization of their expression data.

## Results

### talklr prioritizes ligand-receptor pairs by the specificity of their expression distribution across cell types within a tissue

Ligand-receptor interactions play a critical role in inter-cellular communication. Many single cell RNA-seq studies examine the expression levels of ligand-receptor pairs among different cell types in a tissue to gain biological insights: uncover cell type(s) that are hubs of inter-cellular communications and reveal paracrine/autocrine signaling between cell types^6–11^, many of which are then experimentally validated. For example, ligand-receptor interactions in the NF-κB, FGFR, IGF1R, and JAK pathways were validated as regulators of liver bud development after analyzing the expression patterns of over 2,500 ligand-receptor pairs^6^ among three major cell types. This demonstrated that although ligand-receptor interactions occur at the protein level, analyzing expression patterns of ligand-receptor pairs can lead to experimentally validated new knowledge.

talklr is developed to prioritize ligand-receptor interactions for subsequent validation. We assume that there are n cell types in sufficient proximity to interact with each other (in direct physical contact, or within diffusion limit of soluble ligands), and a gene pair involving a ligand (L) and a receptor (R) mediating the inter-cellular signaling. There are n^2^ possible pairwise cell-cell interactions among the n cell types using the ligand-receptor pair. **Figure 1** gives two toy examples where n=3 (cell types A, B and C). There are three autocrine interactions, where the same cell types express both the ligand and receptor (A-> A, B->B, C->C) and six paracrine interactions, where one cell type expresses the ligand, and a different cell type expresses the receptor (A-> B, A-> C, B-> A, B-> C, C-> A, C-> B). Multiple cell-cell interactions can exist at the same time (e.g., A->B and A-> C), and such one-to-many or many-to-many interactions are indeed known to occur frequently in the literature, as we will show later. Ligand-receptor interactions can be quantified using an interaction score: L_i_*R_j_, the product of expression levels for the ligand in cell type i and the receptor in cell type j^9, 12^ (**Figure 1B**). In talklr, we normalize interaction scores by dividing L_i_*R_j_ with the sum of interaction scores across all possible states, so that normalized interaction scores sum up to 1 (**Figure 1C**).

Researchers often perform single cell RNA-seq of one ^10^ or multiple tissues ^13, 14^ in a baseline condition (often healthy, unperturbed adult condition) to uncover cellular diversity. In this single-condition scenario, talklr compares the observed ligand-receptor interaction score distribution against a reference distribution where both the ligand and receptor are expressed at the same level across all cell types within a tissue (**Figure 1C vs D**). The reference interaction score distribution is simply 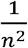for each one of the n^2^ possible interactions, thus exhibiting no cell type specificity (**Figure 1D**). We use the Kullback-Leibler divergence to quantify how the observed distribution of interaction scores differs from the reference distribution (**Figure 1D**). Larger values mean that the observed distribution of interaction scores for a ligand-receptor pair deviate a lot from the evenly distributed reference. In other words, the observed interaction scores are highly concentrated in a few of the n^2^ possible cell-cell interactions. Compared to existing approaches that often only focus on cell type-specific genes^7, 11^ (the expression of a gene is enriched only in one cell type), or ignore cell type-specificity^6, 9^, talklr accommodates all possible expression patterns of the ligand-receptor pair. Cell-type specific expression of individual ligand/receptor genes is sufficient but not necessary for high specificity of the interaction score distribution.

The expression pattern depicted in the left panel of **Figure 1A** fits existing approaches, where both the ligand and the receptor are enriched in only one cell type (not necessarily the same cell type): expressed 3 fold higher in one cell type compared to each of the other two cell types. The KL divergence between the observed and reference interaction score distributions is 0.417. The right panel of **Figure 1A** would be missed by differentially-expressed gene based approaches^7, 11^, because the receptor is highly expressed in two cell types, instead of just one cell type. However, the KL divergence for the right panel is 1.956. This is because the ligand show 10 fold difference between high-vs. low-expression cell types in Figure 1B instead of 3 fold in Figure 1A. More importantly, although the receptor is highly expressed in two cell types instead of one, its expression level in those two cell types is 30 fold higher than the one cell type only weakly expressing it. Thus, the distribution of interaction scores achieved high specificity without individual genes being cell type-specific (**Figure 1B & 1C**). As the number of cell types increases, there are increasingly many ways for the distribution of interaction scores to be highly concentrated in a few possible states (i.e., higher KL divergence), without requiring individual ligand or receptor genes to be uniquely expressed in only one cell type. This is valuable because as the number of cell types increases in complex tissues, it is decreasingly likely that a gene is only enriched in one single cell type. Moreover, talklr also naturally accommodates the common scenario where cell types within a tissue can be related to each other (e.g., CD4+ and CD8+ T cells), because it allows a gene to be highly enriched in a small set of similar cell types.

### talklr identifies ligand-receptor interactions underlying tissue-specific functions

To demonstrate the performance of talklr in the single condition scenario, we applied it on three datasets: bulk RNA-seq of three FACS-sorted cell types (podocytes, endothelial cells, mesangial cells) in the kidney glomerulus^15^, single cell RNA-seq of adult mouse heart from the Tabula Muris project^13^, and bulk RNA-seq of FACS-sorted neurons, astrocytes and microglia from mouse cortex ^16^. We compared the performance of talklr to a commonly used workflow, where both the ligand and receptor are required to be enriched only in one cell type (“DEG pairs”)^7, 11^. In each tissue, we first identified *k* ligand-receptor pairs where both genes are enriched in only one cell type. We also picked the top *k* ligand-receptor pairs with the highest KL divergence (i.e., highly spiked interaction score distribution, “talklr pairs”). We then compared the enrichment of relevant tissue-specific Gene Ontology terms (e.g., kidney development, cardiomyocyte differentiation, and neurogenesis, respectively) in the DEG vs. talklr ligand-receptor gene pairs. talklr pairs showed much higher enrichment for almost all tissue-specific GO biological processes than the DEG-based approach (**Figure 2A**). The number of ligand-receptor pairs uniquely identified by each method make up 23% - 49% of the *k* DEG pairs (**Figure S1A**).

**Figure 2.**
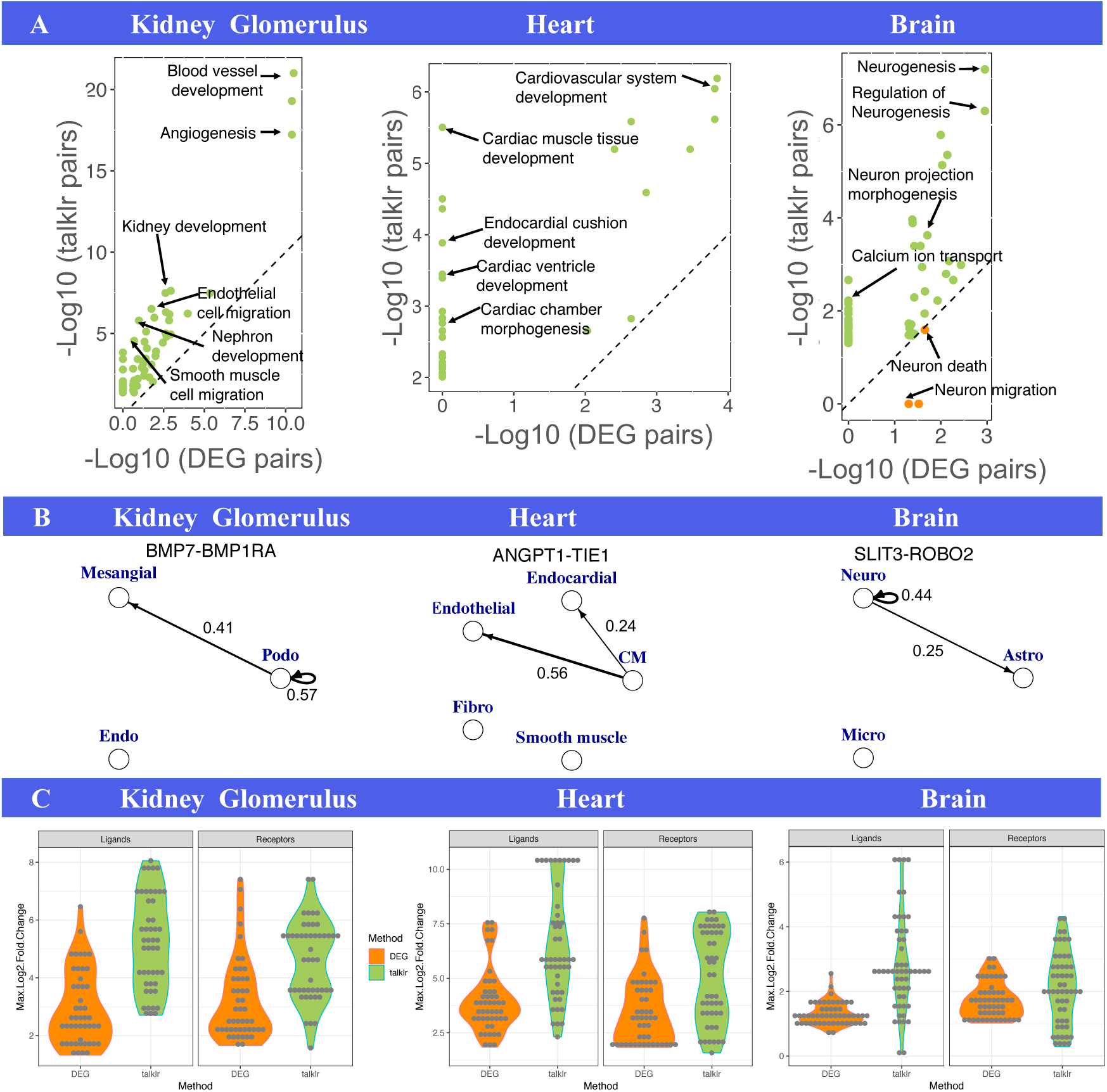
talklr identified ligand-receptor pairs involved in tissue-specific functions. **A.** Scatter plot of GO term enrichment in ligand-receptor pairs selected by the DEG approach (x-axis) and talklr (y-axis) in kidney glomerulus (left), adult heart (middle) and brain (right). A union of significant (FDR<0.05) GO terms among the top 50 GO terms enriched in ligand-receptor pairs selected by either approach are plotted. GO terms with higher enrichment in talklr pairs are colored green; GO terms with higher enrichment in DEG pairs are colored orange. Dashed diagonal line represents equal enrichment. Axes are drawn to scale: longer axes represents larger –log10(FDR), and higher enrichment. Select tissue-specific GO terms are labeled. For fair comparison, we selected the top *k* ligand/receptor pairs from each approach. The number *k* is entirely determined by the number of ligand-receptor pairs where both genes are DEGs. **B.** Select ligand-receptor pairs (Bmp7-Bmpr1a in kidney, Angpt1-Tie1 in heart, Slit3-Robo2 in brain) identified by talklr but missed by DEG approach in three tissues. Each node represents a cell type. Edges go from the cell type expressing the ligand to the cell type expressing the receptor. Numeric labels on edges are normalized interaction scores. To reduce clutter, only large interaction scores are plotted. Note that in all three tissues, talklr pairs are involved in biologically realistic scenarios where one cell type express the ligand and two cell types express the receptor, i.e., one-to-many interactions. **C.** Ligand-receptor pairs selected by talklr have larger expression difference between cell types with highest vs. lowest expression of the ligands and receptors. Each dot is a gene.

talklr uncovered biologically relevant ligand-receptor pairs missed by DEG-based approaches (**Figure 2B**). In the kidney glomerulus, talklr ranked multiple ligand-receptor interactions in the BMP pathway (e.g., Bmp7-Bmpr1a, Bmp7-endoglin) very high. Reassuringly, the BMP pathway is known to play important roles in glomerulus function^17, 18^. These important interactions are missed by the DEG approach because the BMP receptors are expressed by both podocytes and mesangial cells. Similarly, in the heart, the angiopoietin 1 - Tie1 interaction is ranked high by talklr but missed by DEG-based approaches, because the receptor Tie1 is highly expressed in both endocardial and endothelial cells. Cardiomyocyte-derived angiopoietin 1 is known to regulate the function of both the endocardial and endothelial development and function^19, 20^. Finally, in mouse cortex, the Slit3-Robo2 interaction was uniquely discovered by talklr because the receptor, Robo2 is enriched in both neurons and astrocytes. Robo2 is known to be expressed in astrocytes, and the Slit-Robo pathway is known to be important for neuron-glia interaction ^21^. Full talklr results are in table S1-S3.

On the other hand, there are also ligand-receptor pairs identified by DEG approach but not talklr. For example, the Lama1-Sdc4 interaction is uniquely identified by the DEG approach. However, the ligand Lama1 is expressed 4.86 fold higher in podocytes than the other two cell types. This expression difference is much smaller than the BMP ligand-receptor interactions uniquely identified by talklr, where the fold change between expressed and non-expressed cell types for the ligand and the receptor are 215.8 and 51.6, respectively (**Figure S1B**). To demonstrate this systematically, we calculated maximal log2 fold change between the cell types with highest and lowest expression of the ligand and receptor genes. In all three datasets, both ligand and receptor genes identified by talklr have significantly higher maximal log2 fold change than DEG pairs (**Figure 2C**). talklr achieved this because it allows a gene to be highly expressed in more than one cell types, as long as there are large expression differences across cell types (**Figure 1, right panel**). Although DEG is based on fold change, the typical requirement that a gene is enriched in only one cell type tends to be too restrictive. Full DEG pairs results are in table S4-6.

In summary, many known ligand-receptor pairs belong to the situation where one cell type sends the signal, but more than one cell type receive the signal^17–21^. These important one-to-many or many-to-many inter-cellular communication events are missed by existing approaches based on differential expression analysis. Additionally, large expression differences across cell types also mean that talklr is more robust to noise in expression measurements.

### An atlas of tissue-specific ligand receptor interactions across 20 organs and tissues

After demonstrating that talklr performs significantly better than existing approaches at identifying ligand-receptor pairs highly enriched for tissue-specific functions, we applied it on single cell RNA-seq transcriptomes of 20 organs/tissues from the Tabula Muris project^13^. This analysis generated an atlas of ligand-receptor interactions across all major mouse organs, and will be a valuable resource for researchers. In 16 of the 19 tissues, significantly more GO terms showed higher enrichment in ligand-receptor gene pairs selected by talklr than by differential expression-based approach using the binomial test (**Figure 3A**; null hypothesis is that a GO term is equally likely to show higher enrichment in gene pairs selected by either method). In the remaining three tissues (liver, lung, and spleen), more GO terms still showed higher enrichment in talklr pairs than in DEG pairs, but the difference is not statistically significant (**Figure 3A;** FDR are 0.162, 0.163, and 0.33 for liver, lung, and spleen respectively. Full results in table S7). Thus, talklr consistently outperforms DEG-based approaches across a wide range of tissues.

**Figure 3.**
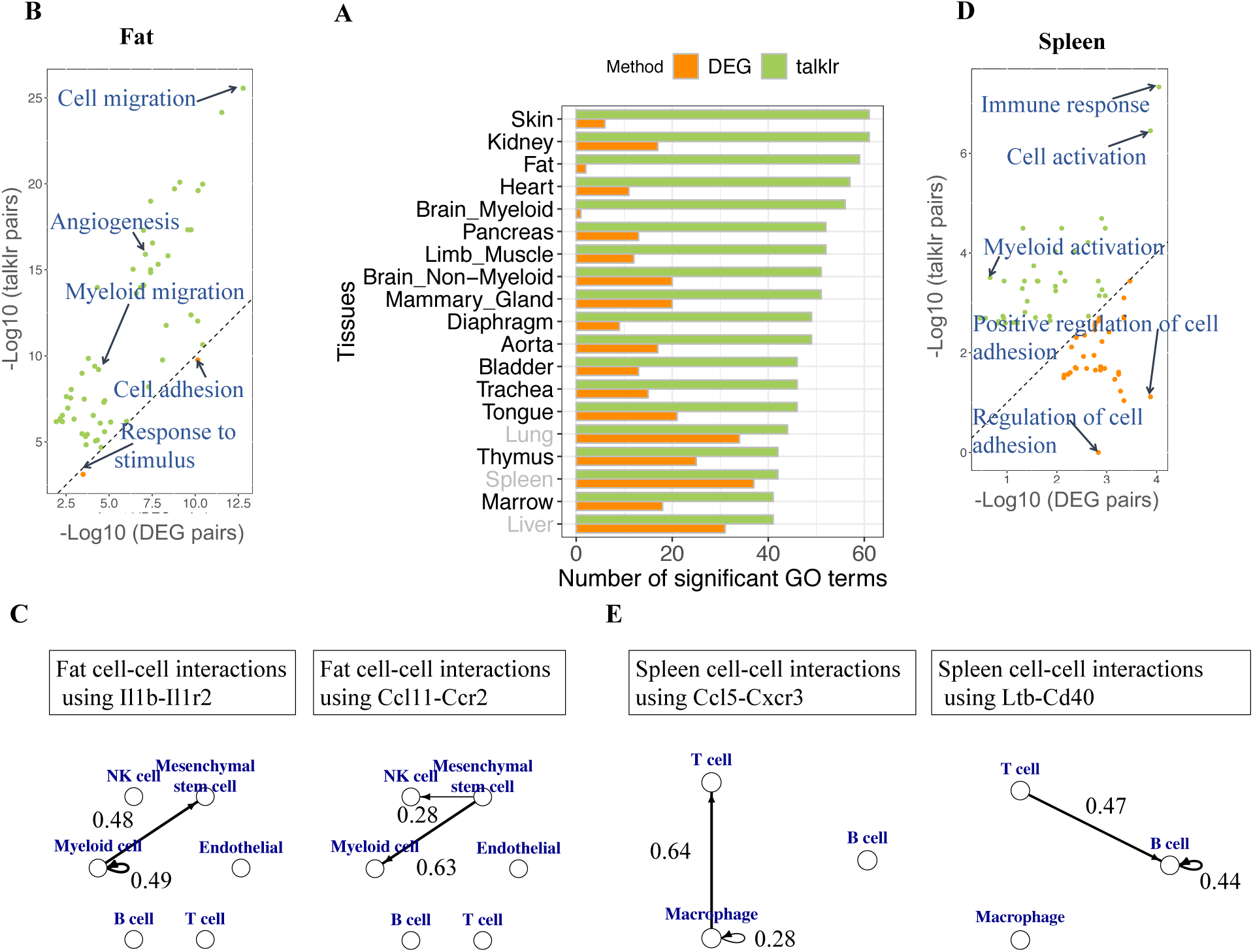
talklr identified tissue-specific ligand-receptor interactions across 20 tissues in the Tabula Muris project **A.** talklr outperforms the DEG-based approach across 19 tissues. A union of GO terms that have FDR<0.05 and ranked among the top 50 most enriched in ligand-receptor pairs selected by either approach are included for analysis. Green bar represents number of GO terms with higher enrichment in talklr pairs; orange bar represents number of GO terms with higher enrichment in DEG pairs. 16 tissues with black font are tissues where there are significantly more GO terms with higher enrichment in talklr pairs than in DEG pairs under binomial test. 3 tissues (lung, spleen, liver) shown in grey are tissues where the number of GO terms with higher enrichment in talklr pairs is still higher than that of DEG pairs, but the difference is not statistically significant. For fair comparison, the same number of ligand/receptor pairs are selected by each approach. **B.** In fat (adipose tissue), all but two GO terms show higher enrichment in talklr pairs than in DEG pairs. **C.** Two examples of cell-cell interactions involving ligand-receptor pairs selected by talklr but missed by DEG approach in fat. Left: myeloid cells express the Il1b ligand (sender), while both MSCs and myeloid cells express the receptor Il1r2 (receivers). Right: MSCs express the ligand Ccl11 (sender), while myeloid cells and to a lesser extent, NK cells, express the receptor Ccr2 (receivers). Normalized interaction scores are labeled on edges. Only interaction scores >0.1 are plotted in order to reduce clutter. **D.** In the spleen, talklr pairs showed higher enrichment in immune response and leukocyte activation GO terms that are known to be spleen relevant. DEG pairs showed higher enrichment for cell adhesion, which is more generic. **E.** Two examples of cell-cell interactions involving ligand-receptor pairs selected by talklr but missed by DEG approach in spleen. Left: macrophages express the Ccl5 ligand (sender), while both macrophages and T cells express the receptor Cxcr3 (receivers). Right: Both T cells and B cells express the ligand Ltb (sender), while only B cells express the receptor Cd40 (receivers). Normalized interaction scores (sum up to 1) are labeled on edges. Interaction scores >0.1 are plotted in order to reduce clutter.

talklr revealed interesting biology in two tissues that we examined in greater detail. In fat (adipose tissue), almost all GO terms showed higher enrichment in talklr pairs than in DEG pairs (**Figure 3B**). Top GO terms with higher enrichment in talklr pairs are myeloid migration and angiogenesis, which are known to regulate adipose tissue function^22, 23^. Many of the ligand-receptor pairs uniquely identified by talklr mediate inter-cellular signaling where one cell type express the ligand, and two cell types express the cognate receptor. For example, adipose mesenchymal stem cells (MSC) express the Il1b ligand, and both NK cells and myeloid cells (macrophages) express the receptor Il1r2. This may be an important mechanism for MSCs to attract both types of immune cells^24^. Similarly, myeloid cells express the Ccl11 ligand, and adipose MSCs and myeloid cells express the receptor Ccr2 (**Figure 3C**). This may mediate the chemotaxis of MSC toward myeloid cells, and mutually influence each other’s phenotypes^25, 26^.

Even when there are similar numbers of GO terms enriched in ligand-receptor pairs selected by both methods, such as in the spleen, GO terms with higher enrichment in talklr pairs are more biologically relevant (**Figure 3D**). The spleen is known to be involved in immune system function, and the ligand-receptor pairs prioritized by talklr showed much higher enrichment for GO terms such as immune response and myeloid activation than DEG pairs. On the other hand, the DEG approach prioritized ligand-receptor pairs involved in cell-cell adhesion and extracellular matrix structure, which may be less relevant for spleen function. For example, Ccl5-Cxcr3 is prioritized by talklr but missed by DEG, because the receptor Cxcr3 is expressed 109.5 normalized UMI counts in T cells and 47.8 in macrophages, and thus did not pass the 3 fold enrichment cutoff. It is well known that Cxcr3 is important in both T cells^27^ and macrophages^28^. T cells and macrophages are known to interact in tumors and other diseases, and Ccl5-Cxcr3 may be important for T cell-macrophage interaction in the spleen (**Figure 3E**). An example where multiple cell types send a signal and one cell type receives it, would be Lymphotoxin B ligand expressed in both T and B cells, while the receptor Cd40 is only expressed in B cells (**Figure 3E**). This interaction may be important for T cell-B cell interactions that are known to occur in the spleen^29^.

In two other immune organs, thymus and bone marrow, there are not only significantly more GO terms with higher enrichment in talklr pairs, these GO terms also belong to immune-related processes. In contrast, DEG pairs showed higher enrichment for cell adhesion (**Figure S2**). This is also observed in another non-immune organ, the tongue, where talklr pairs show higher enrichment for epithelial cell fate commitment and hemidesmosome assembly, known to be important for the tongue^30^, while DEG pairs show higher enrichment for extracellular matrix (ECM) GO terms (**Figure S2**). GO term enrichment scatter plot for all other tissues are shown in **Figure S3**.

In summary, talklr consistently identified ligand-receptor pairs that achieved higher enrichment in tissue-specific biological processes. The atlas of ligand-receptor interactions prioritized by talklr across 20 mammalian tissues will be a valuable resource for researchers to further explore and are available as supplementary data

### talklr identifies disease-perturbed ligand receptor interactions

Single cell transcriptomics is increasingly used to uncover cell type-specific dynamic responses to various perturbations (e.g., diseases, developmental cues, mutations, drug treatments). Uncovering how inter-cellular communications are perturbed in these conditions is a significant challenge. Existing approaches often require one or both of the ligand-receptor genes to be differentially expressed between the baseline and perturbed conditions^8, 31^. We hypothesize that a gene’s statistically significant expression change in one cell type may not be biologically relevant if the gene is highly expressed in other interacting cell types. **Figure 4** showed a common scenario observed in our analysis of many datasets. There is a statistically significant increase of the ligand gene in cell type A, from 5 FPKM in the baseline to 20 FPKM in the perturbed condition. A similar increase occurred for the cognate receptor in cell type A as well (baseline in **Figure 4A vs.** perturbed condition in **4B**). Existing approach would declare this ligand-receptor pair interesting. However, both the ligand and the receptor genes are expressed more than 10 fold higher in cell type C, and their expression in cell type C did not change. Thus, the C->C autocrine interaction is the dominant interaction in both the baseline and perturbed conditions (i.e., distribution of normalized interaction scores did not differ in **Figure 4A vs. B**). Therefore, talklr de-prioritizes this ligand-receptor pair. On the other hand, if the ligand gene also decreased from 200 to 30 FPKM in cell type C, then the distribution of interaction scores changes: both A->C and C->C interactions are important in the perturbed condition. talklr would prioritize this change (**Figure 4 A vs. C**): the KL divergence for Figure 4A vs. 4B and 4A vs. 4C are 0.14 and 1.25, respectively. Thus, talklr not only captures highly relevant ligand-receptor interactions specific to a biological perturbation that are missed by existing DEG-based approach, even for the ligand-receptor pairs identified by both DEG and talklr (e.g., **Figure 4C** vs. **Figure 4A**), talklr provides a quantitative way to rank such ligand-receptor pairs, so that researchers can focus on the top ones.

**Figure 4.**
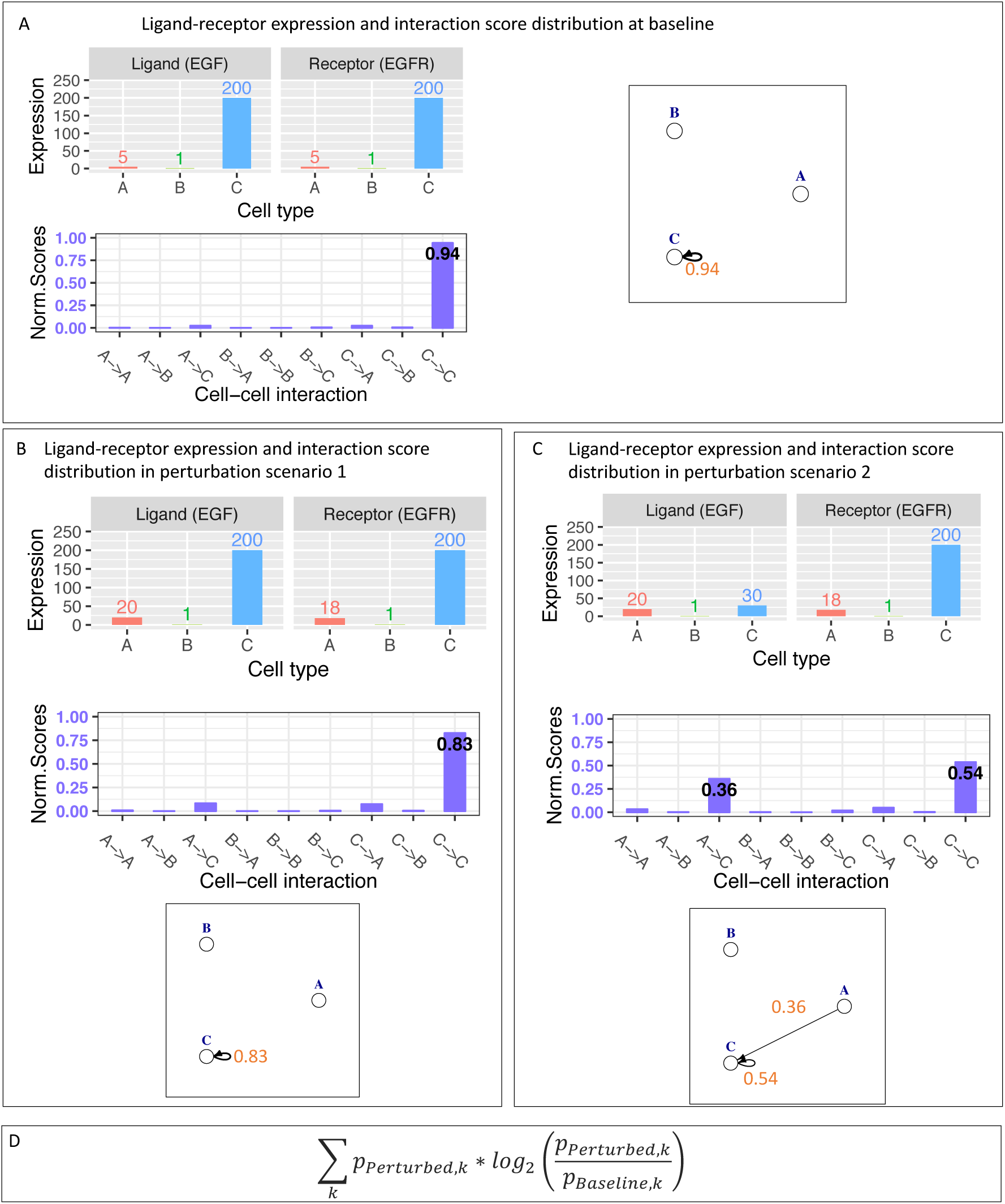
talklr can identify potential ligand-receptor rewiring induced by perturbations in two condition comparison mode. The expression values shown A, B, C are entirely hypothetical. A. Ligand-receptor interaction at baseline. Top left: Ligand/receptor gene expression across three cell types (A, B, C), note that both the ligand and receptor are much more highly expressed in cell type C than in A or B. Bottom left: Distribution of normalized interaction scores. Note that the C->C interaction dominates with a score of 0.94. Right: Network diagram of ligand-receptor interactions among A, B and C. To reduce clutter, only large interaction scores are plotted. B. Ligand-receptor interaction in perturbation scenario 1. Top left: Ligand/receptor gene expression across three cell types (A, B, C). Note that in cell type A, ligand increased to 20 from 5; receptor increased to 18 from 5, compared to the baseline. Note that both the ligand receptor are still much more highly expressed in cell type C than in A or B even after the expression increase. Bottom left: Distribution of normalized interaction scores. Note that the C->C interaction dominates with a score of 0.83. Right: Network diagram of ligand-receptor interactions among A, B and C. To reduce clutter, only large interaction scores are plotted. KL divergence for scenario 1 vs. baseline (Figure 4B vs. A) is 0.14. C. Ligand-receptor interaction in perturbation scenario 2. Top left: Ligand/receptor gene expression across three cell types (A, B, C). Note that in cell type A, ligand increased to 20 from 5; receptor increased to 18 from 5, compared to the baseline. Additionally, in cell type C, ligand expression decreased from 200 to 30. Bottom left: Distribution of normalized interaction scores (sum up to 1). Note that both A-C and C->C interaction have high interaction scores (0.36 and 0.54 respectively). Right: Network diagram of ligand-receptor interactions among A, B and C. Now both cell type A and C send signal, while cell type C receives the signal. Interaction scores >0.1 are plotted. KL divergence for scenario 2 vs. baseline (Figure 4C vs. A) is 1.25, much larger than scenario 1 vs. baseline D. KL divergence compares distribution of interaction scores between perturbed and baseline conditions.

Compared to existing approaches that rely on differential expression of individual genes in individual cell types, talklr takes a holistic view of the expression distribution of both the ligand and the receptor genes across all cell types, and evaluates perturbation-induced expression change of one gene in one cell type in the context of three things: 1. the baseline expression of this gene in other cell types; 2. expression changes of this gene in other cell types and 3). the expression changes of this gene and its cognate ligand/receptor. talklr uses KL divergence to quantify changes in the distribution of interaction scores for all possible cell-cell interactions between two conditions (**Figure 4D**).

To evaluate the performance of talklr, we applied it on three datasets: bulk RNA-seq of FACS-sorted cells from healthy and infarcted heart tissue^32^, immature (new born) vs. mature (postnatal day 56) heart tissue^32^, and single cell RNA-seq of young (2-3 months) and old (21-22 months) mouse brain^33^. We plotted the enrichment p-value for the union of significant GO terms (FDR<0.1) within the top 50 GO terms for DEG-based and talklr approaches in ligand-receptor pairs identified by either approach. Similar to **Figure 2**, most disease-related GO terms show higher enrichment in ligand-receptor pairs prioritized by talklr. For example, myocardial infarction (MI) is known to involve massive cardiomyocyte apoptosis, immune cell infiltration and fibrosis^34^. Ligand-receptor pairs uncovered by talklr showed significant enrichment in cell matrix adhesion, leukocyte migration and apoptotic signaling (not among top 50, but FDR is 0.10 in DEG and 0.006 in talklr), all highly relevant to MI (**Figure 5A, left**). However, ligand-receptor pairs identified based on differential expression analysis did not show significant enrichment in those disease-relevant GO terms. Similarly, in post-natal heart maturation, ligand-receptor pairs uncovered by talklr showed much higher significant enrichment in GO terms related to cardiovascular system development (**Figure 5A, middle**). In brain aging, ligand-receptor pairs identified by talklr showed higher enrichment in biological processes known to be important in brain aging, such as neurogenesis^35^, blood vessel development^36^, and axon development^37^ (**Figure 5A, right**).

**Figure 5.**
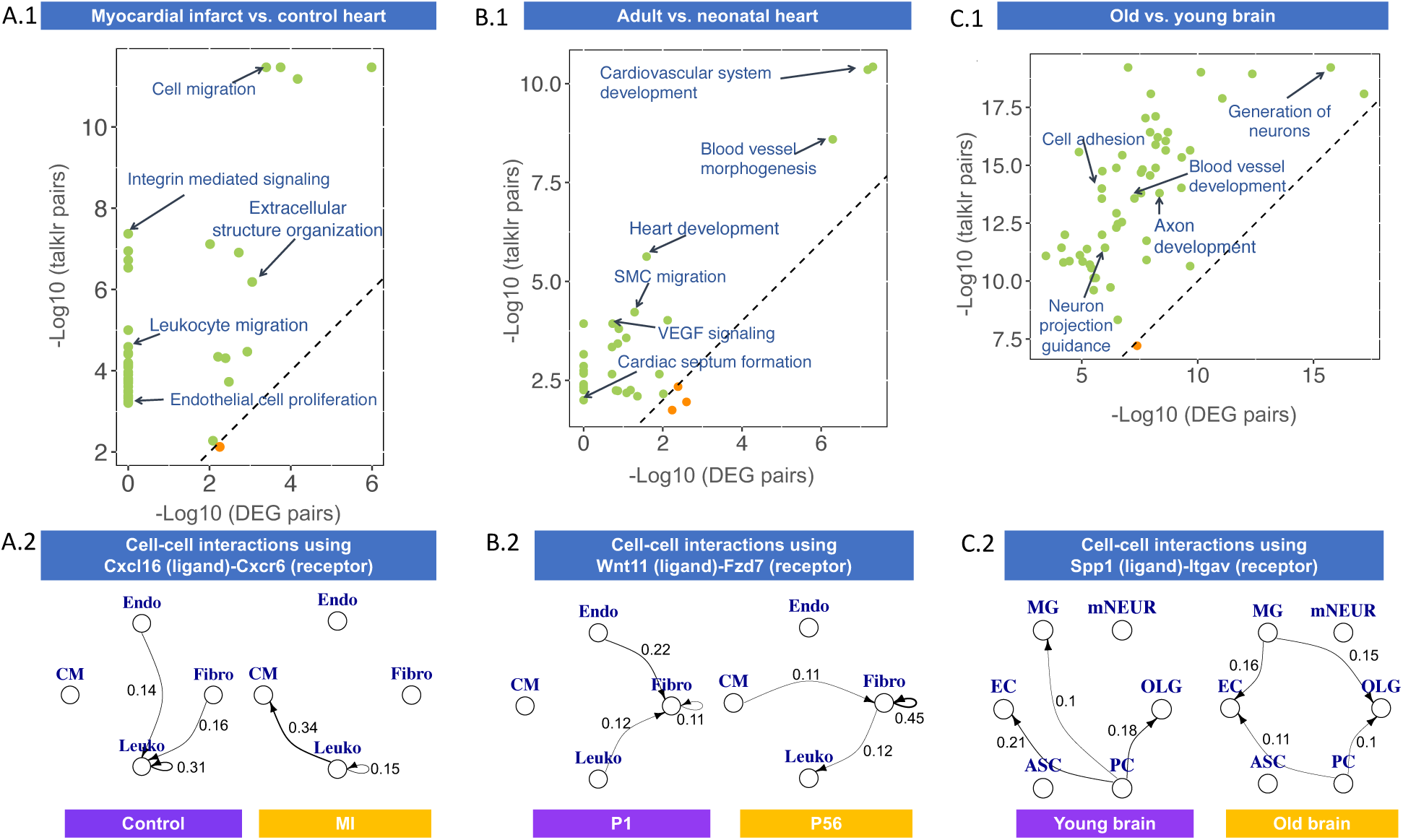
talklr identified ligand-receptor pairs highly relevant to myocardial infarction (left), heart development (middle) and brain aging (right). A. Scatter plot of GO term enrichment in ligand-receptor pairs selected by DEG approach (x-axis) and talklr (y-axis) in myocardial infarction (left), heart development (middle) and brain aging (right). A union of significant (FDR<0.05) GO terms among the top 50 GO terms enriched in ligand-receptor pairs selected by either approach are plotted. GO terms with higher enrichment in talklr pairs are colored green; GO terms with higher enrichment in DEG pairs are colored orange. Dashed diagonal line represents equal enrichment. Axes are drawn to scale: longer axes represent larger –log_10_(FDR), and higher enrichment. Select GO terms known to be relevant for the respective biological processes are labeled. For fair comparison, the same number of ligand/receptor pairs are selected by each approach. B. Select ligand-receptor interactions prioritized by talklr but missed by DEG approach in each biological process. Left: Cxcl16-Cxcr6. In control heart, only leukocytes (Leuko) express the receptor Cxcr6, while leukocytes, and to a lesser extent, fibroblast (Fibro) and endothelial cells (endo), express the ligand Cxcl16. In MI, both cardiomyocytes (CM) and leukocytes express the receptor Cxcr6, and only leukocyte express the ligand Cxcl16. Cxcl16-Cxcr6 has been shown to be causally involved in MI, and reducing it is known to be cardio-protective, possibly reducing cardiomyocyte-leukocyte interaction. Middle: Wnt11-Fzd7. In postnatal day 1 heart, only fibroblasts express the receptor Fzd7; fibroblasts, endothelial cells and leukocytes express the ligand Wnt11. In mature post-natal day 56 heart, both fibroblasts and leukocytes express the Fzd7 receptor, and cardiomyocytes and fibroblasts express the ligand Wnt11. Right: In young brain, Spp1 is expressed mostly by pericytes (PC) and the Itgav receptor is expressed by endothelial cells (EC), microglia (MG) and mature neurons (mNEUR). In old brain, microglia became a major sender of Spp1.

We further examined ligand-receptor pairs identified by each approach in each dataset to reveal new biology (full results are in table S8-10 for talklr and table S11-13 for DEG). In MI, talklr uniquely identified Cxcl16-Cxcr6 interactions as perturbed. Cxcl16-Cxcr6 has been causally linked to MI: disrupting Cxcl16-Cxcr6 signaling is validated to be cardio-protective in mouse models of MI.^38^ Compared to control, the expression of Cxcl16 is more concentrated in leukocytes in MI; and surprisingly, the expression of Cxcr6 shifted from leukocyte to cardiomyocyte (**Figure 5B, left**). Thus, the Cxcl16-Cxcr6 interaction may mediate cardiomyocyte-leukocyte interaction in MI. On the other hand, the Selplg-Itgam interaction is uniquely found by DEG. Although this pair exhibits absolute expression changes, the distribution of interaction scores does not change: it is predominantly leukocyte-leukocyte in both conditions (**Figure S4A**).

In heart development, Wnt11-Fzd7 is uniquely identified by talklr and is known to be important^39^. On the other hand, ligand-receptor pairs identified by DEG often do not have changes in interaction score distribution. For example, the Ccl3-Ccr1 is uniquely selected by DEG approach because Ccl3 expression decreased from 123 transcript per millions (TPM) to 32.5 TPM in leukocytes, and Ccr1 expression decreased from 8.1 TPM to 1.2 TPM in cardiomyocytes. This may not be biologically significant, because Ccr1 expression in leukocytes is 15 fold and 80 fold higher than in cardiomyocytes in immature and mature heart, respectively (**Figure S5B**), and therefore its expression change in cardiomyocyte is not biologically relevant. Indeed, CCR1 is a marker of macrophages (a type of leukocyte), and is not known to be important for cardiomyocyte^40^. This demonstrates that considering the expression change of one gene in one cell type in the context of its expression in other cell types is important.

In brain aging, talklr ranks Spp1 (osteopontin)-Itgav (alpha v integrins) interaction at the top. Spp1 is important in microglia^41^. In young mice, Spp1 is mainly expressed by pericytes. In old mice, microglia is the major sender of Spp1, followed by pericytes. Spp1 is known to promote M2 activation of microglia and may play a protective role^42^. Additionally, the Spp1-Itgav interaction may facilitate the migration and adhesion of microglia toward endothelial cells and oligodendrocytes^43^. On the other hand, the interactions captured by the DEG approach are cases where changes in the expression of the ligand/receptor in one cell type may not be biologically significant because it is more highly expressed in another cell type, and its expression did not change in that high-expressing cell type (e.g., Cd14-Itga4, **Figure S4C**).

Taken together, talklr performs better at identifying perturbation-induced re-distribution of cell-cell ligand-receptor interactions than existing approaches.

## Discussion

Inter-cellular signaling is fundamental to tissue homeostasis and response to perturbations. Recent single cell transcriptomic studies increasingly focus on inferring potential inter-cellular signaling in homeostatic and perturbed conditions. To meet this scientific challenge, we developed a new computational approach called talklr to quantitatively identify important ligand-receptor interactions. We demonstrated that talklr can perform two valuable tasks. First, talklr can uncover ligand-receptor pairs that are enriched for tissue-specific functions (**Figure 2 and 3**), and thus may be important for tissue homeostasis. Second, talklr can identify ligand-receptor rewiring among interacting cell types between two different conditions (e.g., disease vs. healthy, old vs. young, **Figure 4 and 5**). We demonstrated that such ligand-receptor pairs are highly relevant to the biological processes under investigation (**Figure 5**), and may be targeted pharmacologically for disease treatment.

We demonstrated that talklr outperforms existing approaches in these two important tasks. We validated the performance of talklr across many datasets, including the Tabula Muris project, creating an atlas of tissue-specific signaling networks. There are two important reasons underlying the good performance of talklr.

First, talklr considers expression changes in both ligand and receptors, and does so for all cell types simultaneously. Thus, talklr emphasizes large expression differences, is robust to measurement noise, and considers expression changes in one cell type in the context of its baseline expression in other cell types. This is distinct from existing approaches that often rely on cell type-specific or condition-specific differential expression of individual genes in individual cell types, i.e., univariate analysis in both genes and cell types. In other words, talklr prioritize interaction-level specificity, while previous approaches emphasize gene-level specificity.

Second, talklr allows one-to-many or many-to-many cell-cell signaling, while also prioritizing ligand-receptor pairs with cell type-specificity (e.g., expressed 200 TPM in two out of five cell types, <1 TPM in the other three cell types). This enables talklr to capture literature supported, biologically meaningful one-to-many or many-to-many signaling interactions, and it is also a major reason that talklr can discover functional cell-cell interactions missed by existing approaches.

However, we also acknowledge several limitations of talklr. Similar to other approaches, we rely on transcriptomic data, which is not an ideal proxy for ligand and receptor activity. Thus, further experiments are required to verify the functional relevance of the top ligand-receptor pairs prioritized by talklr. Second, we assumed that all cell types within a tissue can interact with each other. This may not be realistic, as certain cell types can be physically too far away to actually interact. However, talklr does allow researchers to specify which cell types interact with each other based on prior knowledge or spatial transcriptomics data. Third, talklr does not consider whether ligand-receptor interactions lead to transcriptional activation of downstream target genes.

In the future, we will extend talklr to 1) utilize transcriptomic technologies that preserves high resolution spatial information, such as spatial transcriptomics^44^, Slide-seq^45^ and seqFISH^46^, as well as computational methods that reconstruct spatial positions of cells without prior information to infer which cell types are actually sufficiently close to interact, either via direct physical contact or through ligand diffusion. 2) to identify ligand-receptor interactions that are perturbed along a continuous trajectory (e.g., inferred pseudo-time trajectory; 3) to further prioritize ligand-receptor pairs by incorporating downstream transcriptional activation of signaling pathways whenever such data is available.

In conclusion, talklr is an information-theory based computational method to prioritize biologically relevant ligand-receptor pairs mediating multi-cellular interactions in homeostatic and perturbed conditions. We believe it will be a valuable addition to single cell RNA-seq analysis workflows.

## Methods

For bulk RNA-seq, the count table from each study was downloaded, and transcript per million (TPM) values were calculated for each gene. For single cell RNA-seq, the UMI tables from each study were downloaded. UMI counts in each cell were normalized to the median counts across all cells. In both types of data, the cell type annotation from the original studies were used. The mean expression level for each gene in each cell type is then calculated. In bulk RNA-seq, only genes expressed above 4 TPM in at least one cell type are considered for further analysis. In single cell RNA-seq, genes with at least 5 normalized UMI counts were further considered.

<u>In single condition mode</u>, talklr prioritizes ligand-receptor pairs that are highly cell type-specific, without requiring that both ligand and receptor genes are enriched only in one cell type.

First, each of the 2400 ligand-receptor pairs, and *n* cell types, L_i_ is the expression of ligand L in cell type i (i=1, 2, 3, …, n). R_j_ is the expression of receptor R in cell type j (j=1, 2, 3, …, n). When *i*=*j*, it is autocrine signaling; when *i* ≠ *J*, it is paracrine signaling. Assuming all cell types can interact with each other, there are n^2^ possible cell-cell interaction pairs.

First, we calculate a vector consisting of raw interaction scores: [L_1_*R_1_, L_1_*R_2_, …, L_2_*R_1_, L_2_*R_2_, …L_i_*R_j_,…,L_n_*R_n_].

Second, we normalize the raw interactions scores to the total sum of interaction scores, S, where 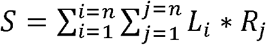. This results in a vector consisting of normalized interaction scores [L_1_*R_1_/S, L_1_*R_2_/S,..L_2_*R_1_/S, L_2_*R_2_/S, …L_i_*R_j_/S, …,L_n_*R_n_/S]

Third, we compared the observed distribution of normalized interaction scores with a background distribution, where the interaction score is the same for each of the n^2^ possible cell-cell interactions [1/n^2^, 1/n^2^, …, 1/n^2^]. The dissimilarity between the observed and background distributions of interaction scores is quantified by KL divergence:

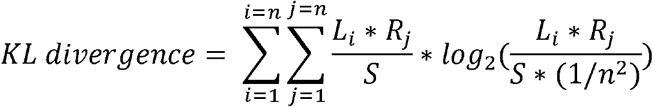

KL divergence is calculated for each of the ligand-receptor pairs where both the ligand and receptor are expressed above a threshold in at least one cell type (expressed, not necessarily enriched). Larger KL divergence means the observed distribution of interaction scores differ more from the background null distribution. In other words, the interaction scores are more concentrated in a few of the n^2^ possible interaction modes, rather than distributed evenly.

<u>In comparison mode</u>, where we aim to uncover potential re-wiring of inter-cellular ligand-receptor interactions between two conditions, we simply replace the reference distribution with the distribution of interaction scores in the baseline condition (e.g., healthy, no treatment):

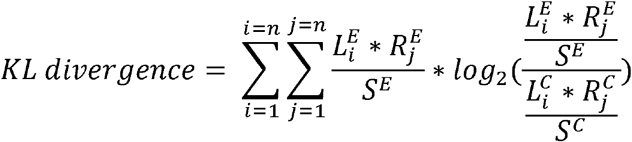

Where 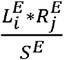 and 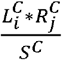 are normalized interaction scores for the perturbed (e.g., disease, treated) and baseline conditions (e.g., healthy, non-treated), respectively.

We compared talklr with the DEG approach, which is commonly used ^7, 8, 31^. In the single condition mode, only ligand-receptor pairs where both the ligand and receptor are enriched in only one cell type are considered (3 fold higher in one cell type compared to each of the other cell types). This approach thus only considers one-to-one cell-cell interaction. To compare talklr and DEG approach in the single condition mode, we first identified a number, *k*, of ligand-receptor pairs where both ligand and receptor are expressed in a cell type-specific manner. Then we selected the top *k* ligand-receptor pairs with highest KL divergence calculated using talklr. We then performed Gene Ontology enrichment analysis for the genes in the top *k* DEG and talklr pairs. We then plotted false discovery rate of GO terms that satisfy two criteria: 1) among the top 50 most enriched GO terms in either approach and 2) FDR<0.05 in either approach. We then plotted these GO terms and examined which approach achieved higher enrichment p-value for each of the GO terms.

We quantified whether talklr performs better than the DEG approach by checking whether there are significantly more GO terms having smaller p-values in the talklr approach compared to the DEG approach using the binomial distribution. The null hypothesis is that the two approach perform equally well, p=0.5, i.e., a GO term is equally likely to have higher enrichment in either approach.

All computational analysis was performed using the R programming language. The R package *igraph* was used to plot networks. *pheatmap* was used to make heatmaps. *ggplot2* was used to make scatter plots and violin plots. *topGO* package was used to perform Gene Ontology enrichment.

### Data availability

This study used both bulk RNA-seq of FACS-sorted pure cell populations, as well as single cell RNA-seq data. Accession number and additional information about them are listed below:

**Table 1.** List of public datasets re-analyzed in this study.

| Accession number | Technology | Description |
| --- | --- | --- |
| GSE129788 | scRNA-seq | Mouse brain aging |
| GSE75246 | Bulk RNA-seq | Mouse Alzheimer's model (PS2APP) |
| GSE95755 | Bulk RNA-seq | Mouse model of myocardial infarction and maturation |
| GSE123179 | Bulk RNA-seq | Mouse kidney glomerulus |
| GSE109774 | scRNA-seq | Mouse Tabula Muris project (20 tissues) |

### Code availability

The talklr R package is available at Github at https://github.com/yuliangwang/talklr.

The talklr interactive website is available https://yuliangwang.shinyapps.io/talklr/. This website does not require any programming experience. Users can upload their expression tables, download results and plot ligand-receptor wiring diagrams.

### Supplementary tables and data

Supplementary tables 1-3: ligand-receptor pairs identified by talklr for kidney, heart and brain.

Supplementary tables 4-6: ligand-receptor pairs identified using differentially expressed genes (DEG) for kidney, heart and brain.

Supplementary table 7: binomial test results comparison the number of GO terms with higher enrichment in ligand-receptor pairs identified by talklr or DEG approaches Supplementary table 8-10: ligand-receptor pairs identified by talklr for myocardial infarction, heart development, and brain aging.

Supplementary table 11-13: ligand-receptor pairs identified by DEG for myocardial infarction, heart development, and brain aging.

talklr results for 20 Tabula Muris tissues are provides as a zipped folder of Excel spreadsheets.

## Acknowledgement

This project is supported by the Institute for Stem Cell & Regenerative Medicine at University of Washington. We thank our colleagues Larry Ruzzo, Carol Ware, Diana Eng, Karol Bomsztyk and Peiyun Zhou for critical inputs of the manuscript.

## Figures and figure legends

**Figure S1.**
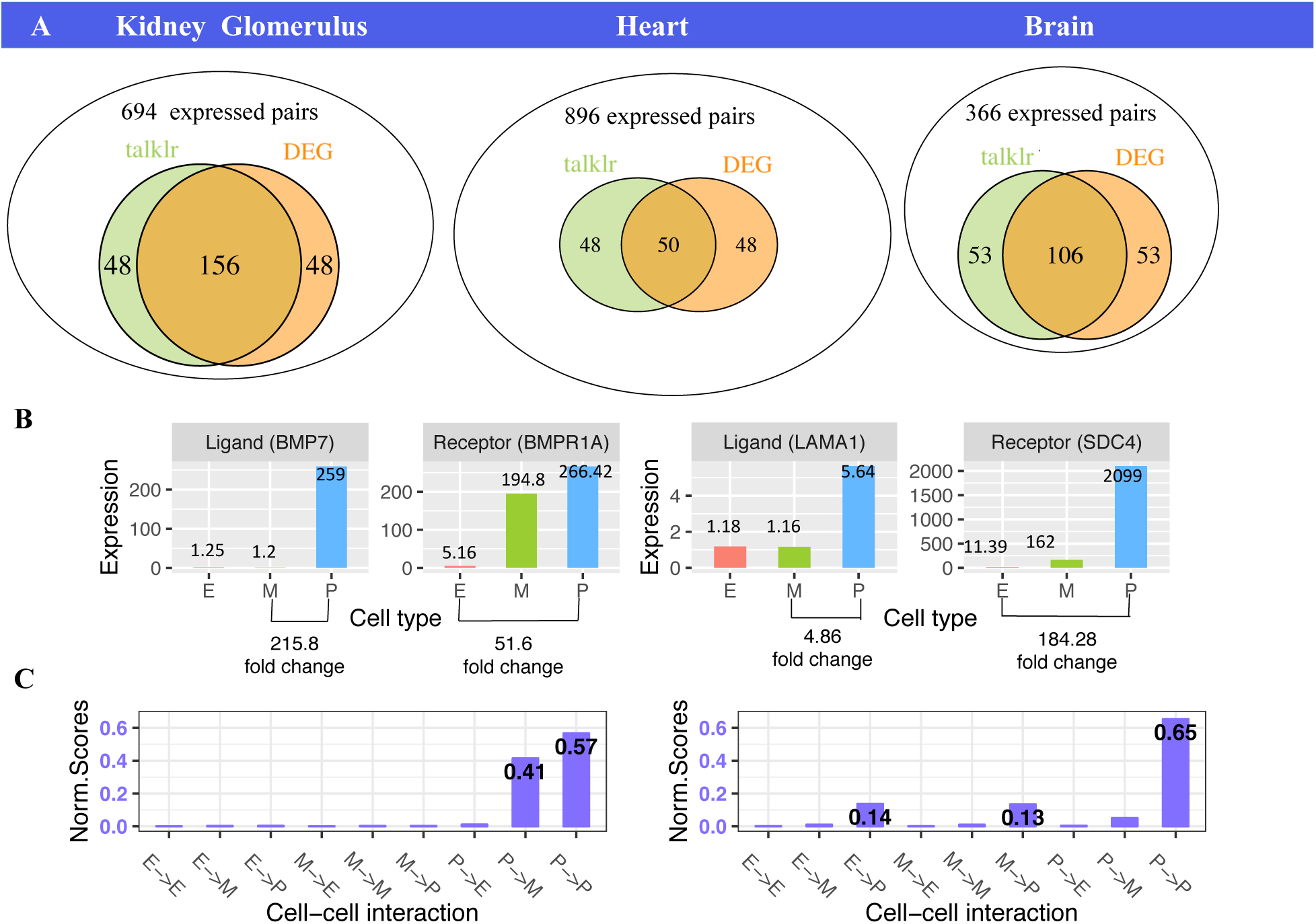
A. Venn diagram of ligand-receptor pairs identified by talklr and DEG pairs. For fair comparison in GO enrichment, the same number of ligand-receptor pairs are selected, and entirely determined by the number of ligand-receptor pairs where both genes are DEGs. In kidney, 23.5% ligand-receptor pairs are unique to each method; in heart, 49% are unique; in brain, 33.3% are unique to each method. B. Ligand-receptor pairs uniquely identified by DEG approach have smaller expression difference. E: endothelial cells; M: mesangial cells; P: podocytes. Bmp7-Bmpr1a is uniquely identified by talklr. The fold change between highest expressing cell type (podocytes) and lowest expressing cell type (mesangial cells) are 215.8 and 51.6 fold for the ligand and receptor, respectively. On the other hand, in the Lama1-Sdc4 pair uniquely identified by DEG, only the receptor Sdc4 has large expression difference between cell types (184 fold), the ligand Lama1 has only 5 fold expression difference. C. Distribution of normalized interaction scores for Bmp7-Bmpr1a and Lama1-Sdc4 Bmp7-Bmpr1a has higher KL divergence than Lama1-Sdc4 (2.02 vs. 1.58).

**Figure S2.**
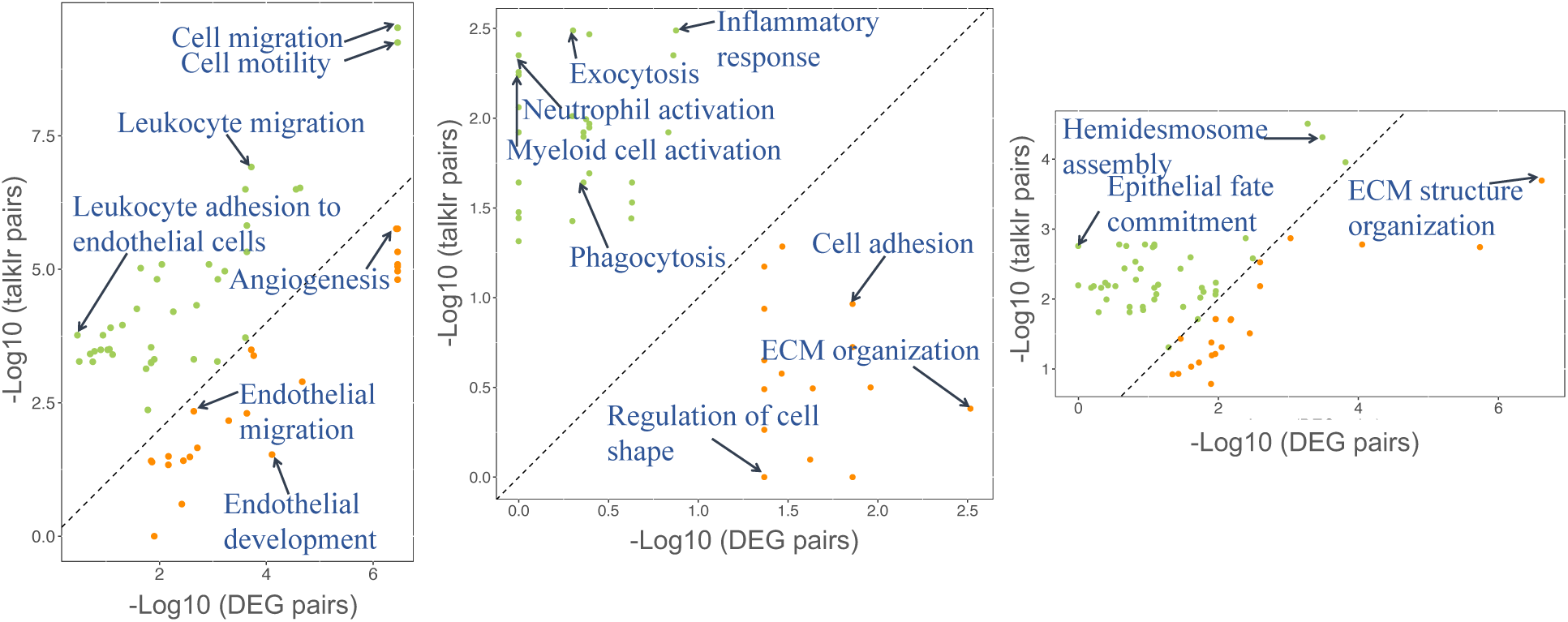
Ligand-receptor pairs prioritized by talklr show higher enrichment in tissue-relevant GO terms than DEG pairs. Scatter plot of GO term enrichment in ligand-receptor pairs selected by DEG approach (x-axis) and talklr (y-axis) in thymus (left), bone marrow (middle) and tongue (right) tissues from the Tabula Muris project. A union of significant (FDR<0.05) GO terms among the top 50 GO terms enriched in ligand-receptor pairs selected by either approach are plotted. GO terms with higher enrichment in talklr pairs are colored green; GO terms with higher enrichment in DEG pairs are colored orange. Dashed diagonal line represents equal enrichment. Axes are drawn to scale: longer axes represent larger –log_10_(FDR), and higher enrichment. For fair comparison, the same number of ligand/receptor pairs are selected by each approach. Select tissue-specific GO terms are labeled.

**Figure S3.**
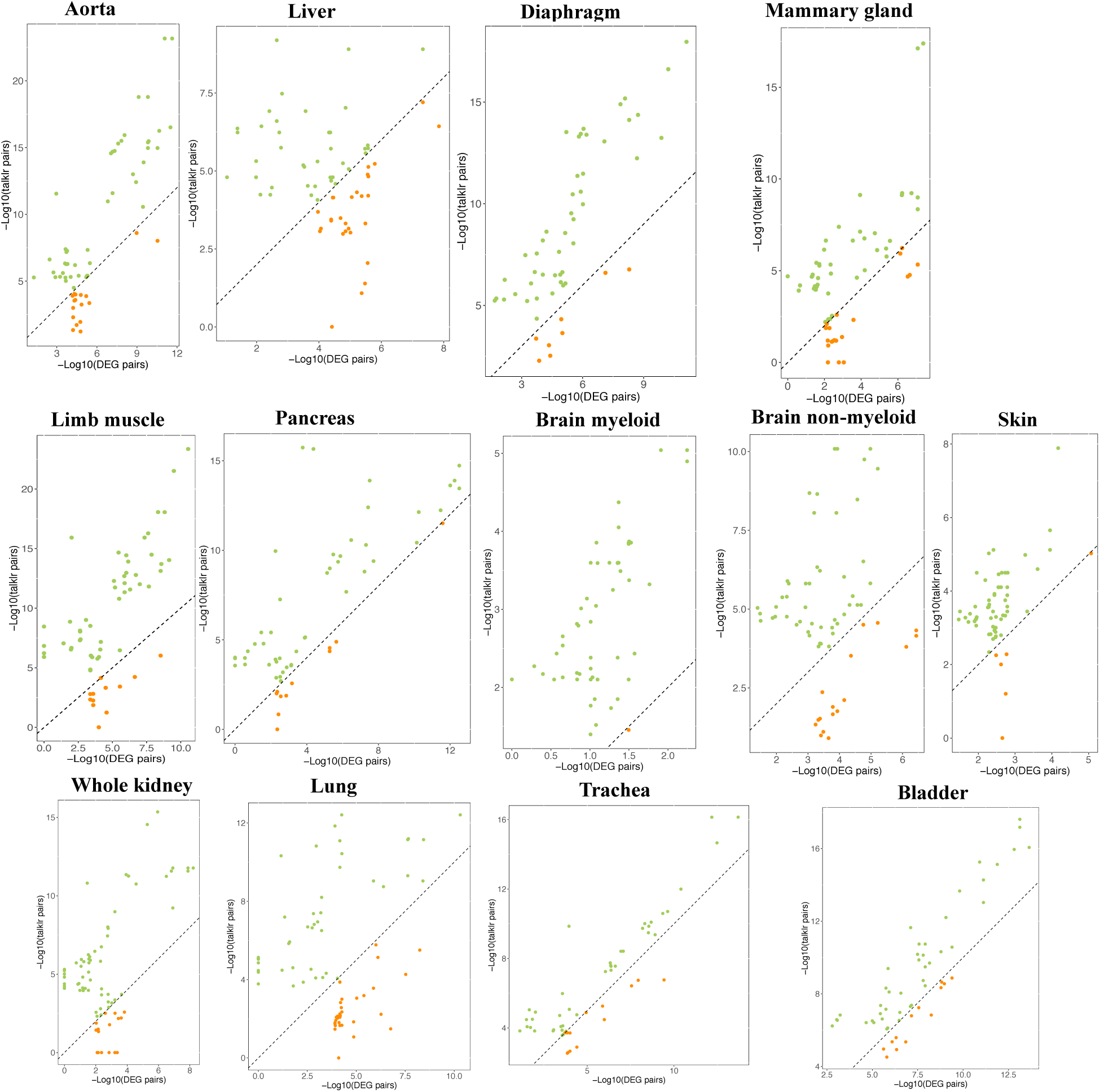
Scatter plot of GO term enrichment in ligand-receptor pairs selected by DEG approach (x-axis) and talklr (y-axis) in 13 tissues from the Tabula Muris project (additional 6 in Figure 2, Figure 3 and Figure S2). A union of significant (FDR<0.05) GO terms among the top 50 GO terms enriched in ligand-receptor pairs selected by either approach are plotted. GO terms with higher enrichment in talklr pairs are colored green; GO terms with higher enrichment in DEG pairs are colored orange. Dashed diagonal line represents equal enrichment. Axes are drawn to scale: longer axes represent larger –log_10_(FDR), and higher enrichment. For fair comparison, the same number of ligand/receptor pairs are selected by each approach. In all tissues, there are more green dots than orange dots.

**Figure S4.**
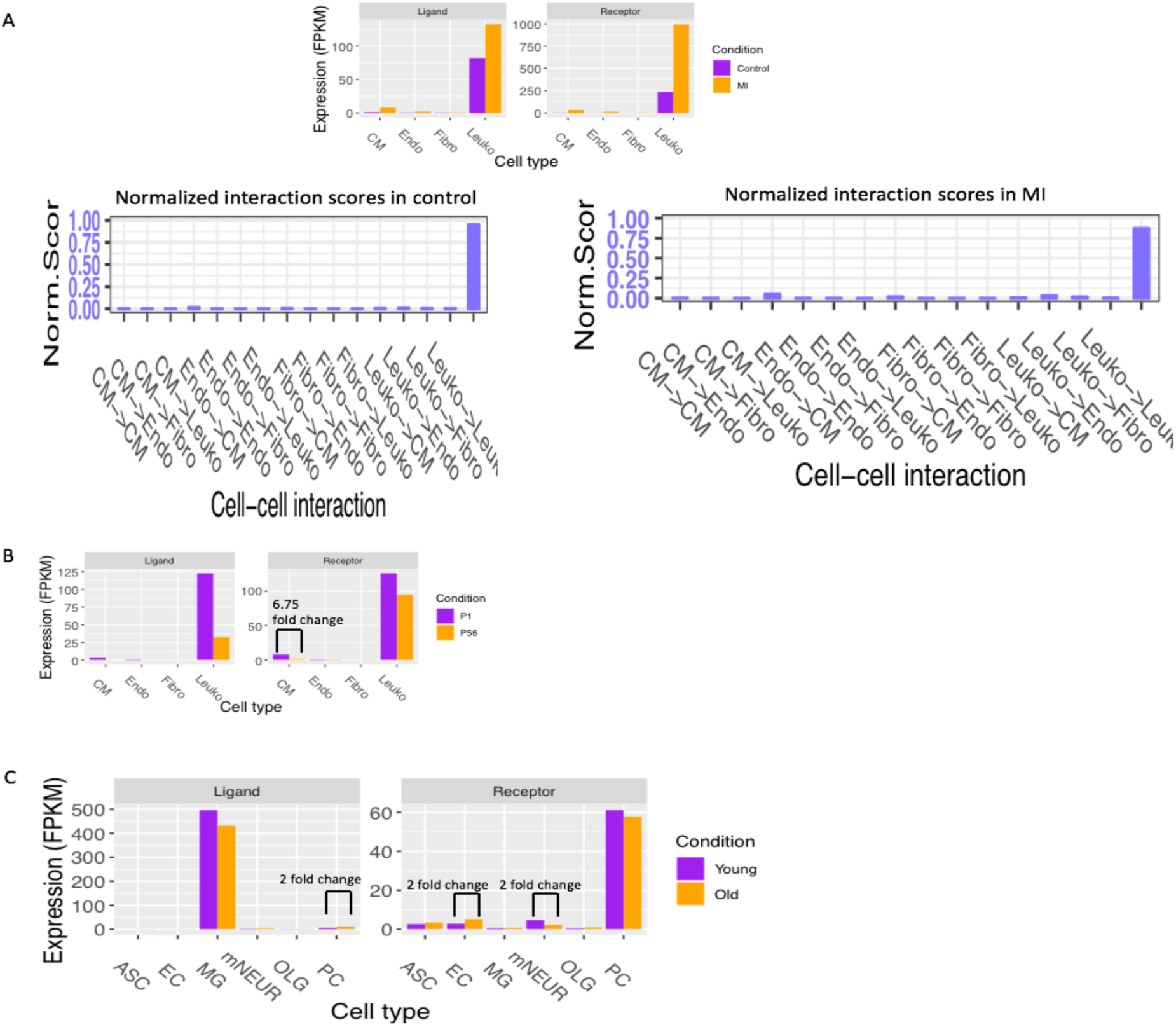
A. The Selplg-Itgam interaction is uniquely identified by DEG approach as perturbed between myocardial infarction (MI) and control heart. However, in both control and MI, both the ligand and the receptor are predominantly expressed by leukocytes, shown by bar plot at the top. As a result, in both control and MI, the leukocyte-leukocyte interaction is the dominant interaction for this ligand-receptor pair, as shown by distribution of interaction scores (bottom). B. The Ccl3-Ccr1 interaction is uniquely identified by DEG approach, because the ligand showed >3 fold reduction in leukocyte, and the receptor showed >3 fold change in cardiomyocyte in heart maturation. However, in both conditions, both the ligand and the receptor are predominantly expressed by leukocytes, shown by bar plot at the top. As a result, in both P1 and P56, the leukocyte-leukocyte interaction is the dominant interaction for this ligand-receptor pair. Moreover, the 3 fold reduction of Ccr1 in CM may not be biologically relevant as Ccr1 is mainly expressed in leukocytes. C. The Cd14-Itga4 interaction is uniquely identified by the DEG approach, because both the ligand and receptor showed >2 fold change in at least one cell type. However, these expression changes may not be biologically significant because they are more highly expressed in other cell types, where expression levels do not change between young and old conditions.

